# Interfacial Tension Modulates Viscous Microfluidic Droplet Generation

**DOI:** 10.64898/2025.12.11.693026

**Authors:** Aarthi Namasivayam, Christopher J. Halbrook, Elliot E. Hui

## Abstract

Mammalian cell culture in nanoliter droplets of extracellular matrix has recently attracted interest as a platform for high throughput drug screening on 3D tissue models. Microfluidic droplet generation using basement membrane extract, such as Matrigel or Cultrex, is complicated by the fact that the viscosity is orders of magnitude higher than that of water. Consequently, conditions that produce quality water droplet generation can fail with basement membrane extract due to the higher capillary number. Here, a parametric study using a T-junction device demonstrates that higher viscosity can be abrogated by lower interfacial tension, which is modulated by surfactant concentration.

## INTRODUCTION

Organoid culture models are rapidly being adopted for drug screening applications. Miniaturization is strongly advantageous in this context, achieving higher throughput at lower cost and reducing the often significant expansion time required to prepare primary patient-specific organoid cultures at sufficient volume.^1,2^ Droplet microfluidics has been successfully employed to generate microencapsulated models consisting of three-dimensional spheroids or organoids in microfluidic droplets of various extracellular matrix materials.^2–4^ Organoids are most commonly grown in basement membrane extract (BME) such as Matrigel (Corning) or Cultrex (Bio-Techne). While automated liquid handling can dispense BME volumes down to 3 µL with 18% coefficient of variance (CV),^5^ this remains an order of magnitude greater than the BME volumes that can be achieved by droplet microfluidics.^2^

Basement membrane extract (BME) has complex rheological properties, displaying strong temperature dependence, shear thinning, and orders of magnitude higher viscosity than is typical in droplet microfluidics,^6^ thus complicating the physics of droplet generation. Using a T-junction droplet generator, we found that conditions that work well for generation of water droplets can fail when trying to form BME droplets (Fig. 1). Droplet formation in a T-junction is a function of the capillary number *Ca = Uμ/γ*, where *U* (m/s) is the flow velocity, *μ* (kg/ms) is the dynamic viscosity, and *γ* (kg/s^2^) is the interfacial tension between the dispersed and continuous phases. At lower *Ca*, monodisperse droplets are generated in the squeezing and dripping regimes, whereas higher *Ca* produces less uniform droplets in the jetting regime or no droplets in the threading regime (Fig. 1B).^7–9^ This behavior has previously been described as a question of whether the interfacial velocity *γ/μ* is sufficiently greater than the flow velocity *U* to allow droplets to snap off cleanly.^7^ Under our experimental conditions (∼12 ºC, 10-100 s^-1^ shear rate), we estimate BME viscosity to be 1-10 Pa s, or 3-4 orders of magnitude higher than that of water.^6,10,11^ The resulting increase in Ca is consistent with the difference in droplet formation between water and BME seen in Fig. 1. Here, we examine the remaining parameters of the Ca equation, flow velocity and interfacial tension,^12,13^ aiming to counteract the high viscosity of BME.

**Figure 1.**
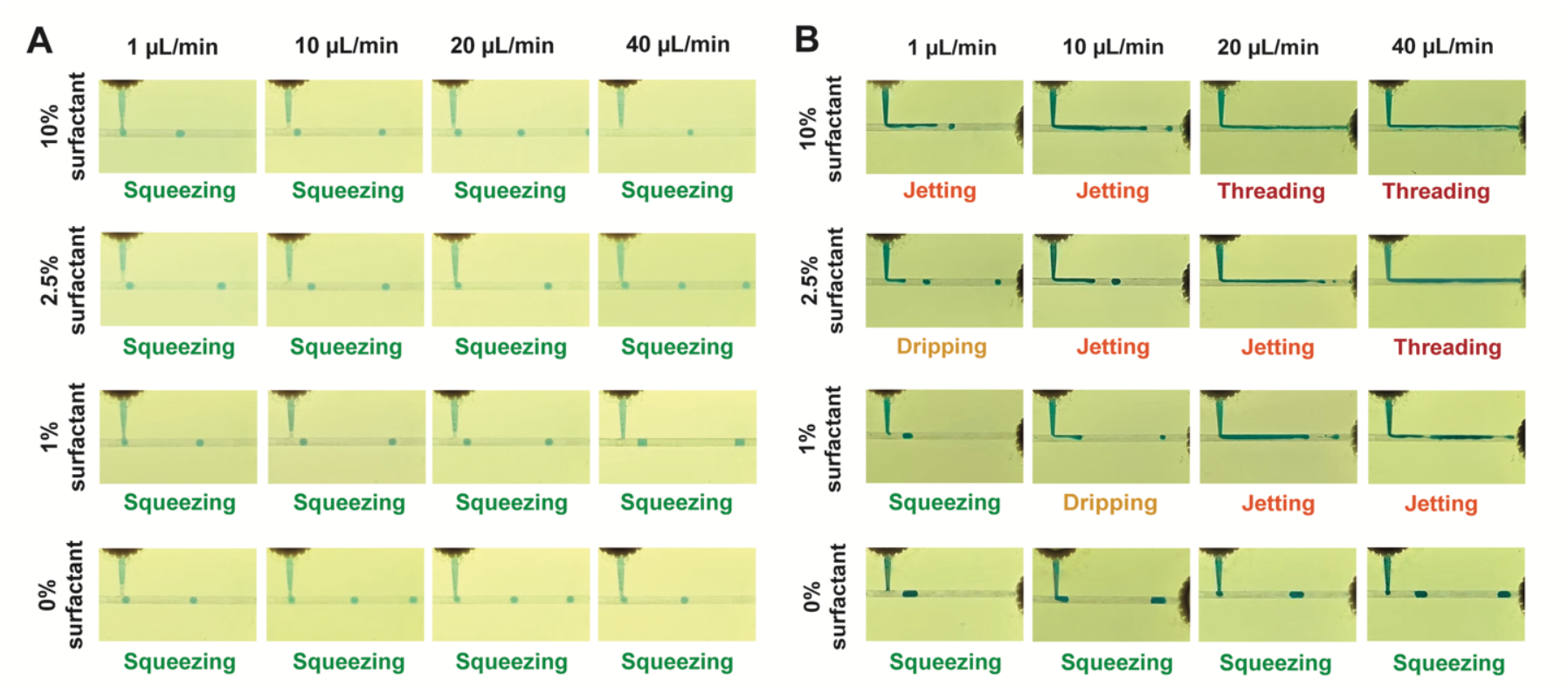
Droplet generation as function of surfactant concentration and flow velocity. (A) Water droplet generation remains constant across a range of surfactant concentrations and flow velocities. (B) Cultrex BME droplet generation is highly dependent on surfactant concentration and flow velocity. Dispersed phase flow rates are given, and a 1:10 flow ratio of dispersed phase to continuous phase was employed in all cases.

## MATERIALS AND METHODS

Droplet emulsions were prepared using a continuous phase of Novec HFE 7500 Engineered Fluid (Applied Thermal Fluids) with Krytox 157 FSH oil (PFPE) surfactant (Grainger), and a dispersed phase of Cultrex BME Type R-1 (Bio-Techne) or deionized water. A microfluidic T-junction was 3D printed with HTL resin on a microArch S240 (Boston Micro Fabrication) printer. The main channel was 1 mm wide, 0.5 mm deep, and 45 mm long. The dispersed phase channel was 1 mm wide, tapering to 0.5 mm at the intersection with the main channel, 0.25 mm deep, and 8.5 mm long. Two syringe pumps (NE-300, New Era Pump Systems) were connected to the microfluidic chip via IDEX NanoPorts (Cole-Parmer). An ice pack was placed around the BME syringe to prevent premature gelation.

## RESULTS AND DISCUSSION

In order to maintain low Ca for quality droplet generation in the face of high BME viscosity, flow velocity can be reduced or interfacial tension can be increased. In this work, we studied the effects of both parameters on droplet generation. Specifically, we modulated interfacial tension by adjusting surfactant concentration,^13^ as surfactants serve to reduce the interfacial tension between liquids. Concurrently, we studied reductions in total flow velocity while maintaining a 1:10 flow ratio of the dispersed and continuous phases. As shown in Fig. 1, both adjustments produced substantial changes in BME droplet generation while minimally affecting water droplet generation.

BME droplets were successfully generated by employing low flow rate or low surfactant concentration. The highest rates of droplet production were achieved without surfactant. Droplets were generated at a maximum of 1.3 Hz with a dispersed phase flow rate of 40 µL/min. Lower droplet frequencies were produced at lower flow rates: 0.44 Hz at 20 µL/min, 0.2 Hz at 10 µL/min, and 0.06 Hz at 1 µL/min. Without surfactant, droplet generation exhibited only a slight decrease in droplet volume as flow rate and capillary number increased, consistent with the squeezing regime (Fig. 1B). Instead, droplet size was determined primarily by the flow ratio of the dispersed and continuous phases, again consistent with the expected behavior in the squeezing regime.^8^ Smaller 317 nL droplets (14.3% CV, n=68) were produced with a 1:15 flow ratio, increasing to 609 nL (10.6% CV, n=50) with a 1:10 flow ratio, and further increasing to 720 nL (16.2% CV, n=30) with a 1:5 flow ratio (Fig. 2A).

**Figure 2.**
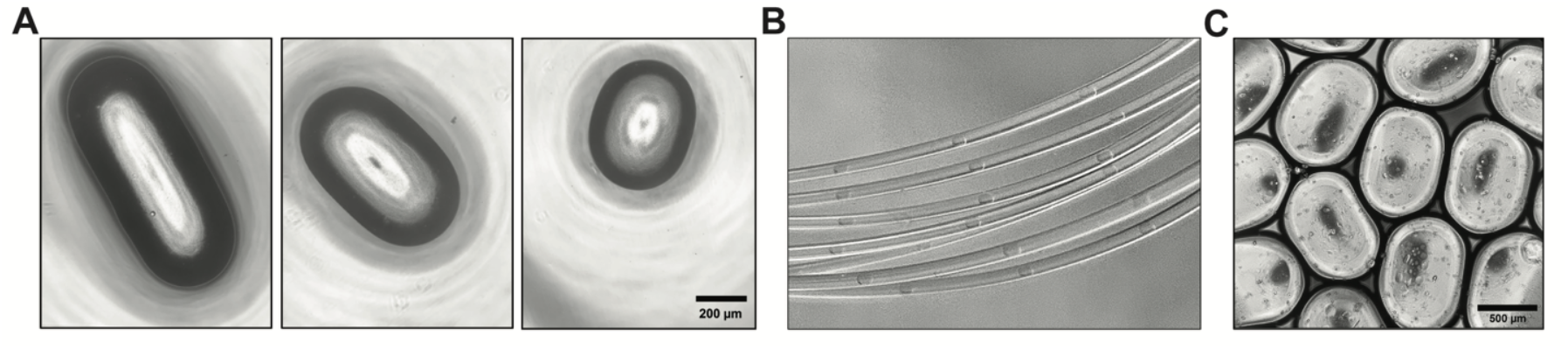
Control of droplet size and gelation. (A) Droplet size varies as a function of the dispersed:continuous flow ratio: 1:5, 1:10, or 1:15 (left to right).(B) Evenly spaced droplets of Cultrex BME collected in the outlet tubing for incubation. (C) Gelation is complete after incubating for 30 min, allowing BME droplets to be dispensed and aggregated without coalescence.

While eliminating surfactant was important for establishing low capillary number in the face of high BME viscosity, surfactants maintain the stability of emulsions and prevent droplet coalescence. To avoid droplet fusion, BME plugs were collected single-file in a spool of 1/32” ID PTFE tubing (Fig. 2B) and incubated for 30 minutes (37 ºC, 5% CO_2_) to gel the BME.^14^ During incubation, the tubing was plugged on both ends by ∼1 cm cubes of PDMS punctured to a depth of ∼5 mm with a 1-mm biopsy punch. Following gelation, the BME beads could be aggregated without fusion (Fig. 2C). Droplet volumes were measured after gelation and dispensing by measuring length and width from images and estimating height from the tubing diameter.

## CONCLUSION

600 nL droplets of Cultrex basement membrane extract were produced at 1 Hz with 10% CV in a microfluidic T-junction. Despite the high viscosity of basement membrane extract, low capillary number was achieved by eliminating surfactant to increase interfacial tension, allowing droplet generation to operate in the robust squeezing regime. We anticipate that our strategy of engineering capillary number through interfacial tension will be instructive for those wishing to generate droplet emulsions with high viscosity fluids.

## ACKNOWLEDGEMENTS

This work was supported by the National Science Foundation through the Center for Advanced Design and Manufacturing of Integrated Microfluidics (IIP-1841509). Additional support was provided by NIH R01GM134418 and the Chao Family Comprehensive Cancer Center (P30CA062203).

## REFERENCES

1 S. Bose, H. Clevers, and X. Shen, “Promises and challenges of organoid-guided precision medicine,” Med 2(9), 1011–1026 (2021).

2 S. Ding, C. Hsu, Z. Wang, N.R. Natesh, R. Millen, M. Negrete, N. Giroux, G.O. Rivera, A. Dohlman, S. Bose, T. Rotstein, K. Spiller, A. Yeung, Z. Sun, C. Jiang, R. Xi, B. Wilkin, P.M. Randon, I. Williamson, D.A. Nelson, D. Delubac, S. Oh, G. Rupprecht, J. Isaacs, J. Jia, C. Chen, J.P. Shen, S. Kopetz, S. McCall, A. Smith, N. Gjorevski, A.-C. Walz, S. Antonia, E. Marrer-Berger, H. Clevers, D. Hsu, and X. Shen, “Patient-derived micro-organospheres enable clinical precision oncology,” Cell Stem Cell 29(6), 905-917.e6 (2022).

3 L. Yu, M.C.W. Chen, and K.C. Cheung, “Droplet-based microfluidic system for multicellular tumor spheroid formation and anticancer drug testing,” Lab. Chip 10(18), 2424–2432 (2010).

4 W. Jiang, M. Li, Z. Chen, and K. W. Leong, “Cell-laden microfluidic microgels for tissue regeneration,” (2016).

5 H. Sherman, J. Shyu, and H. Hung, “Automation of Forskolin-induced Swelling Assay of Human Intestinal Organoids,” (2020).

6 K.I.W. Kane, E. Lucumi Moreno, C.M. Lehr, S. Hachi, R. Dannert, R. Sanctuary, C. Wagner, R.M.T. Fleming, and J. Baller, “Determination of the rheological properties of Matrigel for optimum seeding conditions in microfluidic cell cultures,” AIP Adv. 8(12), 125332 (2018).

7 J.D. Tice, A.D. Lyon, and R.F. Ismagilov, “Effects of viscosity on droplet formation and mixing in microfluidic channels,” Anal. Chim. Acta, (2004).

8 M.D. Menech, P. Garstecki, F. Jousse, and H.A. Stone, “Transition from squeezing to dripping in a microfluidic T-shaped junction,” J. Fluid Mech. 595, 141–161 (2008).

9 X. Chen, T. Glawdel, N. Cui, and C.L. Ren, “Model of droplet generation in flow focusing generators operating in the squeezing regime,” Microfluid. Nanofluidics 18(5–6), 1341–1353 (2015).

10 T. Thorsen, R.W. Roberts, F.H. Arnold, and S.R. Quake, “Dynamic pattern formation in a vesicle-generating microfluidic device,” Phys. Rev. Lett. 86(18), 4163–4166 (2001).

11 S.N. López-Carranza, M. Jenny, and C. Nouar, “Pipe flow of shear-thinning fluids,” Comptes Rendus Mécanique 340(8), 602–618 (2012).

12 T. Dinh, and T. Cubaud, “Role of Interfacial Tension on Viscous Multiphase Flows in Coaxial Microfluidic Channels,” Langmuir 37(24), 7420–7429 (2021).

13 L. Peng, M. Yang, S. Guo, W. Liu, and X. Zhao, “The effect of interfacial tension on droplet formation in flow-focusing microfluidic device,” Biomed. Microdevices 13(3), 559–564 (2011).

14 S. Jiang, H. Zhao, W. Zhang, J. Wang, Y. Liu, Y. Cao, H. Zheng, Z. Hu, S. Wang, Y. Zhu, W. Wang, S. Cui, P.E. Lobie, L. Huang, and S. Ma, “An Automated Organoid Platform with Inter-organoid Homogeneity and Inter-patient Heterogeneity,” Cell Rep. Med. 1(9), 100161 (2020).

